# Brain responses to different types of salience in antipsychotic naïve first episode psychosis: An fMRI study

**DOI:** 10.1101/263020

**Authors:** Franziska Knolle, Anna O Ermakova, Azucena Justicia, Paul C Fletcher, Nico Bunzeck, Emrah Düzel, Graham K Murray

## Abstract

Abnormal salience processing has been suggested to contribute to the formation of positive psychotic symptoms in schizophrenia and related conditions. Previous research utilising reward learning or anticipation paradigms has demonstrated cortical and subcortical abnormalities in people with psychosis, specifically in the prefrontal cortex, the dopaminergic midbrain and the striatum. In these paradigms, reward prediction errors attribute motivational salience to stimuli. However, little is known about possible abnormalities across different forms of salience processing in psychosis patients, and whether any such abnormalities involve the dopaminergic midbrain. The aim of our study was, therefore, to investigate possible alterations in psychosis in neural activity in response to various forms of salience: novelty, negative emotion, targetness (task-driven salience) and rareness/deviance. We studied 14 antipsychotic naïve participants with first episode psychosis, and 37 healthy volunteers. During fMRI scanning, participants performed a visual oddball task containing these four forms of salience. Psychosis patients showed abnormally reduced signalling in the substantia nigra/ventral tegmental area (SN/VTA) for novelty, negative emotional salience and targetness; reduced striatal and occipital (lingual gyrus) signalling to novelty and negative emotional salience, reduced signalling in the amygdala, anterior cingulate cortex and parahippocamal gyrus to negative emotional salience, and reduced cerebellar signalling to novelty and negative emotional salience. Our results indicate alterations of several forms of salience processing in patients with psychosis in the midbrain SN/VTA, with additional subcortical and cortical regions also showing alterations in salience signalling, the exact pattern of alterations depending on the form of salience in question.

## Introduction

Salience is a property that enables a stimulus to attract attention, and to drive cognition and behaviour. It can be described as a product of matched/mismatched stimulus features and internal, driving factors of an individual, such as goals, beliefs and experiences at a particular point in time. Salience is a multifaceted concept ^1^, including different dimensions, such as reward and threat prediction, prediction error, novelty, emotional salience or rareness/deviance. The literature well describes the role of dopamine (DA) for reward prediction error ^2,3^, with neural signals originating in the substantia nigra/ventral tegmental area (SN/VTA)^4^. However, DA neuron firing is not exclusive to reward prediction error, but has also been reported in response to non-rewarding unexpected events, such as aversive or alerting ^5^, as well as novel events ^6^, suggesting that DA release, at least in some contexts, reflects general salience ^7,8^.

In psychosis, abnormal salience processing secondary to dysregulation of the dopaminergic system – described as the ‘aberrant salience’ hypothesis of psychosis ^1,9,10^ – has been linked to the formation and maintenance of psychotic symptoms ^11–13^. It has been suggested that aberrant salience attribution in psychosis is caused by faulty DA signalling in the striatum, possibly driven by dysregulation from the prefrontal cortex (PFC) and hippocampus ^14^. In psychosis, there is increased synthesis and release of DA in the striatum, which is present even at the prodromal stages of the disease ^15,16^. Several studies reported reduced midbrain, striatal, and/or cortical processing of reward prediction errors ^17–20^ and non-reward related prediction errors in psychosis^21^. In our recent work, we documented meso-cortico-striatal prediction error deficits, involving midbrain, striatum and right lateral frontal cortex in medicated psychosis patients at different stages ^19,21^ and in unmedicated first episode psychosis patients and patients at clinical risk for developing psychosis ^22^. Another study in people at clinical risk of psychosis showed a relation between the striatal reward prediction signal and psychotic symptoms ^23^.

Novel events activate DA neurons even in the absence of reward, which is associated with increased attention, memory and goal-directed behaviour ^5^. Together with the fact that novelty exploration engages the areas of the brain involved in appetitive reinforcement learning (i.e. dopaminergic midbrain areas, striatum, medial prefrontal cortex) ^24,25^, novelty may be intrinsically rewarding, irrespective of the choice outcome, and can provide a ‘bonus’ for exploration ^26^. A recent study by Schott and colleagues ^27^ reported alterations in a fronto-limbic novelty processing network in unmedicated (not necessarily first episode) patients with acute psychosis. However, it is unclear whether novelty processing is disrupted in key dopaminergic regions for salience processing, such as the SN/VTA.

Emotional events are also salient, capture attention, enhance memory and modify behavioural responses ^24^. Presynaptic DA levels in the amygdala and SN/VTA predict brain activity in response to emotional salience ^28^. Schizophrenia and first episode psychosis patients have problems processing emotions, especially in the context of facial recognition ^29^. In a PET study, Taylor and colleagues ^30^ showed impaired neural processing in the ventral striatum in response to emotional salient events in chronic and acute psychosis patients. However, results regarding processing alterations in the amygdala were unclear. Furthermore, it is unknown whether processing of the dopaminergic SN/VTA is altered in psychosis in response to emotional salience.

Various studies suggest that SN/VTA neurons respond to a general form of salience (see reviews ^7,8^), sometimes referred to as ‘physical salience’ or ‘alerting’ salience ^2,31^, which is triggered by unexpected sensory events including surprise, attention, arousal, or novelty. If dopaminergic signalling is generally compromised in psychosis, it then follows that there should be overlapping patterns of abnormal activation to various forms of salient stimuli in the dopaminergic midbrain and associated target regions in psychosis patients. Under an alternative account, salience processing may still be generally impaired in psychosis, but this may be secondary to dysfunction of diverse neural systems. In the current study, we, therefore, investigated brain responses in the SN/VTA and other target areas to four types of salience ^25^ – stimulus novelty, negative emotional salience, rareness/deviance (or ‘contextual deviance’), and targetness (task-driven attentional salience) – in patients with early psychosis and healthy volunteers. By focussing on early psychosis, we can avoid confounds of exposure to dopaminergic medications and other effects of chronic illness. We used a fMRI paradigm ^25^ that previously was shown to significantly activate parts of the midbrain, amygdala and striatum to various forms of salience, and which importantly provides an baseline condition (a neutral oddball event) that is matched in frequency with other conditions of interest.

Based on the potentially general role in salience signalling of DA neurons in the SN/VTA and the ‘aberrant salience’ hypothesis of psychosis, we hypothesised that psychosis patients demonstrate altered SN/VTA and striatal responses to novelty, negative emotional salience and targetness. Furthermore, we predicted to find group differences in the prefrontal cortex in response to novelty, in the amygdala in response to emotional salience, and in the hippocampus in responses to all forms of salience.

## Methods

### Subjects

We recruited 14 antipsychotic naive individuals with first-episode psychosis and active psychotic symptoms from the Cambridge, early intervention service for psychosis, CAMEO. Other inclusion criteria were as follows: age 16–35 years, meeting ICD-10 criteria for a schizophrenia spectrum disorder (ICD-10: F20, F22, F23, F25, F28, F29) or affective psychosis (ICD-10: F30.2, F31.2, F32.3). Age, gender and handedness matched healthy volunteers (n=37) were recruited as control subjects. We recruited a higher number of controls than patients in order to improve statistical power, given the challenges in recruiting antipsychotic naïve individuals with active psychotic symptoms (yet still well enough to tolerate the MRI procedure) in their first episode of illness. Demographic and clinical characteristics of those participants included in the final analysis are presented in Table 1 and 2. None of the healthy volunteers reported any personal or family history of severe neurological, psychiatric or medical disorders. All participants had normal or corrected-to-normal vision, and had no contraindications to MRI scanning. At the time of the study, none of the participants were taking antipsychotic medication or had drug or alcohol dependence.

**Table 1:** Sample characteristics for healthy controls, and patients with first-episode psychosis (FEP).

| Variable | Controls<br>(n=34) |  | FEP<br>(n=13) |  | Group Statistics |  |  |
| --- | --- | --- | --- | --- | --- | --- | --- |
|  | Mean | SD | Mean | SD | T | df | p |
| Age (years) | 22.85 | 3.3 | 23.85 | 6.3 | 0.21 | 47 | 0.48 |
| Gender (male/female) | 18/16 |  | 8/6 |  | 0.45 | 47 | 0.61 |
| Handedness<br>(right/left) | 27/7 |  | 9/3<br>(1 missing) |  | 0.06 | 46 | 0.76 |
| **IQ** | 119.4 | 18.2 | 105.1 | 16.2 | 3.03 | 47 | **0.02** |
| Level of education | 2.47 | 0.8 | 2.38 | 1.1 | 0.51 | 47 | 0.77 |
| Mother's level of<br>education | 2.36 | 1.1 | 2.73 | 1.3 | 0.8 | 43 | 0.36 |
| Smoking (yes/no) | 0.32 | 0.5 | 0.54 | 0.5 | 1.61 | 46 | 0.18 |
| **Alcohol** | 2.6 | 0.7 | 1.54 | 1.5 | 3.34 | 46 | **0.002** |
| Cannabis | 0.85 | 0.8 | 1.38 | 1.6 | 2.08 | 45 | 0.12 |
| Hallucinogens | 0.21 | 0.5 | 0.46 | 0.7 | 1.32 | 46 | 0.15 |
| Stimulants | 0.41 | 0.7 | 0.77 | 0.8 | 1.87 | 46 | 0.15 |
| Depressants | 0.06 | 0.2 | 0.23 | 0.6 | 1.33 | 46 | 0.16 |

Education was measured on a 5-point scale (from no education to higher university degree). Intelligence was measured with the Culture Fair Intelligence Test. Smoking: 0=non-smoker, 1=smoker. Substance use was measured on a 5-point scale (from 0=never used to 5=daily user). **Bold:** significant differences.

**Table 2:** Clinical assessment of the participants.

| Variable | Controls (n=34) |  | FEP (n=13) |  | Group statistics |  |  |
| --- | --- | --- | --- | --- | --- | --- | --- |
|  | Mean | SD | Mean | SD | t | df | p |
| **PANSS positive** | 0.97 | 0.2 | 2.2 | 0.7 | 10.1 | 45 | **<0.001** |
| **P1** | 0.97 | 0.2 | 3.2 | 1.6 | 7.86 | 45 | **<0.001** |
| **P2** | 0.97 | 0.2 | 1.3 | 0.9 | 2.22 | 45 | **<0.001** |
| **P3** | 0.97 | 0.2 | 3.4 | 1.6 | 9.06 | 45 | **<0.001** |
| SANS score | 0.09 | 0.3 | 0.3 | 0.6 | 1.6 | 45 | 0.116 |
| **BDI** | 3.4 | 3.9 | 27.8 | 10.7 | 11.1 | 41 | **<0.001** |

FEP, first episode psychosis patients. BDI, Beck's Depression Inventory. PANSS, Positive and negative syndrome scale for schizophrenia. P1, delusions, P2 conceptual disorganisation (thought disorder), P3, hallucinatory behaviour. SANS, Scale for Assessment of Negative Symptoms. **Bold:** significant differences.

Before scanning, each of the participants underwent a general interview and clinical assessment using the Positive and Negative Symptom Scale (PANSS) ^32^, the Scale for the Assessment of Negative Symptoms (SANS) ^33^ and the Global Assessment of Functioning (GAF) ^34^. The Beck Depression Inventory (BDI) ^35^ was used to assess depressive symptoms during the last two weeks. IQ was estimated using the Culture Fair Intelligence Test ^36^.

The study was approved by the Cambridgeshire 3 National Health Service research ethics committee. All participants supplied written informed consent after they had read a complete description of the study.

### Novelty task

We used a visual oddball paradigm ^37^ in order to investigate four types of salience, which were novelty, negative emotional salience, targetness and rareness/deviance. Participants were presented with a series of greyscale images of faces and outdoor scenes. 66.6% of those were used as ‘standard’ images, which were of neutral emotional valence. The four types of rare or contextually deviant events were randomly intermixed with these; each occurred with a probability of 8.3%. These deviant events were: neutral stimuli that required a motor response (‘target oddball’); stimuli that evoked a negative emotional response (‘emotional oddball’, angry face or image of car crash); novel stimuli (‘novel oddball’, different neutral images that appear only once); and neutral stimuli (‘neutral oddball’, neutral image of face or scene) (Figure 1). All participants completed four blocks with 60 trials each, resulting in a total of 240 trials (160 standard trials, and 20 oddball trials each of target, neutral, emotional and novel stimuli). The task contained 50% faces and 50% outdoor scenes, this allowed prevention of category-specific habituation. These categories were chosen instead of abstract images to make stimulus exploration biologically relevant. Participants were introduced to the target stimulus prior to the experimental session for 4.5s, and they were required to make a simple button press with their right index finger in response to each of its subsequent appearances during the experiment within the fMRI-scanner. No motor responses were associated with any of the other stimulus types.

**Figure 1.**
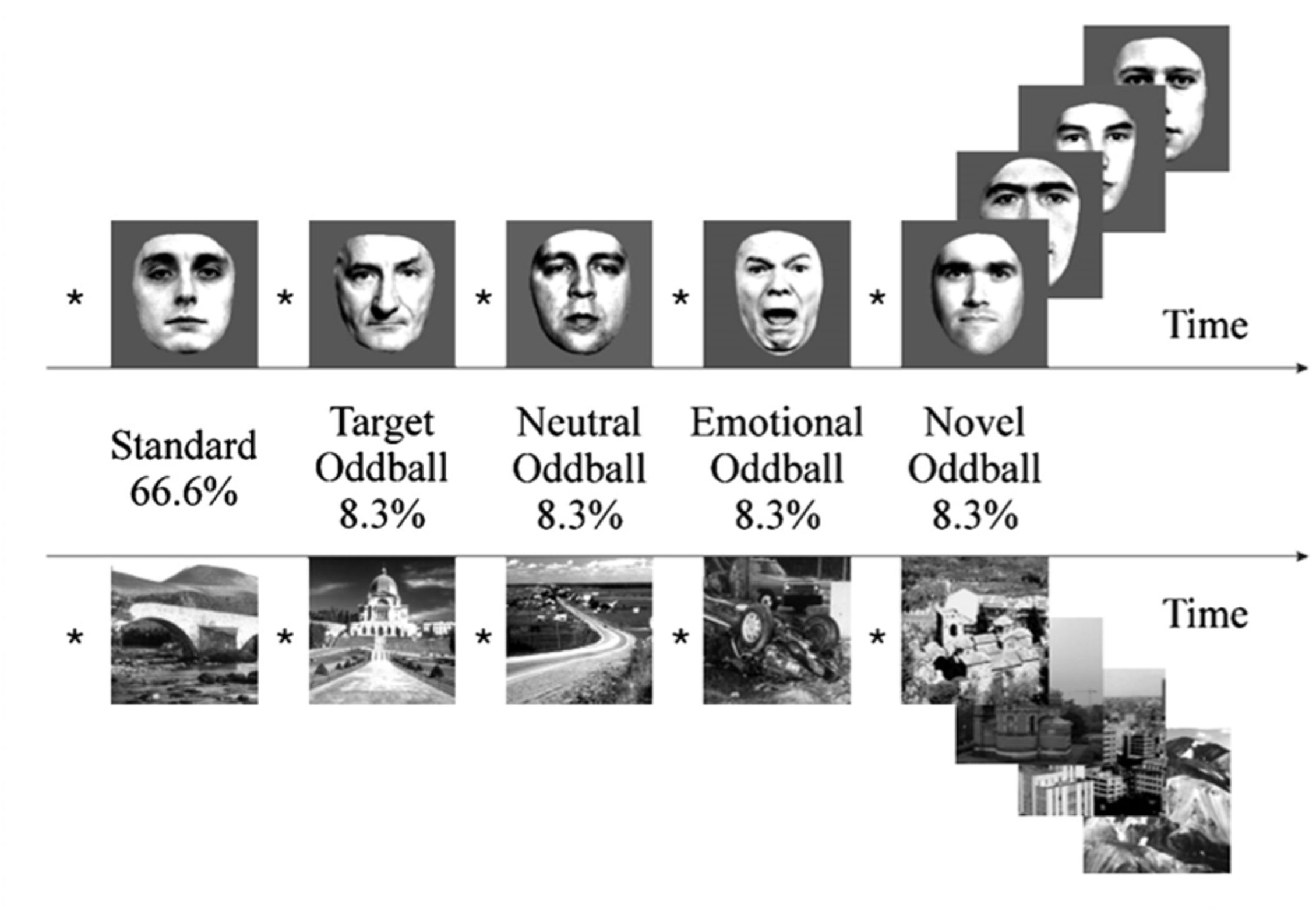
Visual oddball paradigm. Participants are presented with a series of greyscale images of faces and outdoor scenes. 66.6% of those were ‘standard’ images. The remaining 33.4% consisted of four types of rare or contextually deviant events, which were randomly intermixed with the standard stimuli; each occurred with a probability of 8.3%. These deviant events were: neutral stimuli that required a motor response (‘target oddball’); stimuli that evoked a negative emotional response (‘emotional oddball’, angry face or image of car crash); novel stimuli (‘novel oddball’, different neutral images that appear only once); and neutral stimuli (‘neutral oddball’, neutral image of face or scene).

During the fMRI-experiment, the pictures were presented for 500ms followed by a white fixation cross on a grey background (grey value=127) using an inter-stimulus interval (ISI) of 2.7s. ISI was jittered with ±300ms (uniformly distributed). The order of stimuli was optimised for efficiency with regard to estimating stimulus-related haemodynamic responses.

All of the stimuli were taken from Bunzeck and Düzel ^25^. The scalp hair and ears of faces were removed artificially; the outdoor scenes did not include faces. All pictures were grey scaled and normalised to a mean grey value of 127 (SD 75). The pictures were projected on to the centre of a screen, and the participants watched them through a mirror mounted on the head coil, subtending a visual angle of about 8°. The negative emotional scene depicted a negatively rated car accident (without any people). The contrast between stimuli allowed us to examine brain responses to the pure stimulus novelty (‘novel’ vs. ‘neutral’), targetness (‘target’ vs. ‘neutral’), negative emotional valence (‘emotional’ vs. ‘neutral’) and rareness/deviance per se (‘neutral’ vs. ‘standard’) (Figure 1). By contrasting the specific salient oddball events with the neutral oddball events, and including a separate, additional, “rareness/deviance” contrast (that corresponds to a classical pure oddball effect), we can differentiate activation to the various forms of salience under study. Irrespective of whether participants were left or right handed, they used their right hand to press the buttons on the button box for the target picture.

### Behaviour analysis

An analysis of variance (ANOVA) was used to investigate group differences in pressing the buttons in response to the target stimuli and assessing reaction times. All runs in which participants missed more than five button presses were excluded. Behavioural data were analysed using SPSS 21 (IBM Corp.).

### fMRI data acquisition and analysis

A Siemens Magnetom Trio Tim syngo MR B17 operating at 3 T was used to collect imaging data. Gradient-echo echo-planar T2*-weighted images depicting BOLD contrast were acquired from 35 non-contiguous oblique axial plane slices of 2mm thickness to minimise signal drop-out in the ventral regions. We did not retrieve images of the whole brain; the superior part of the cortex was not imaged. The relaxation time was 1620ms, echo time was 30ms, flip angle was 65°, in-plane resolution was 3×3×3mm, matrix size was 64×64, field of view was 192×192mm, and bandwidth was 2442Hz/px. A total of 437 volumes per participant were acquired (35 slices each of 2mm thickness, inter-slice gap 1mm). The first five volumes were discarded to allow for T1 equilibration effects.

The data were analysed using FSL software (FMRIB’s Software Library, www.fmrib.ox.ac.uk/fsl) version five. Participants’ data (first-level analysis) were processed using the FMRI Expert Analysis Tool (FEAT). Functional images were realigned, motion corrected (MCFLIRT^38^) and spatially smoothed with a 4mm full-width half-maximum Gaussian kernel. A high-pass filter was applied (120s cut-off). All images were registered to the whole-brain echo-planar image (EPI) (i.e., functional image with the whole-brain field of view), and then to the structural image of the corresponding participant (MPRAGE) and normalised to an MNI template, using linear registration with FSL FLIRT. The five explanatory variables (EVs) that we used were the onset times of the standard, target, emotional, novel and neutral pictures. They were modelled as 1s events and convolved with a canonical double-gamma response function. We added a temporal derivative to the model to take into account possible variations in the haemodynamic response function. To capture residual movement-related artefacts, six covariates were used as regressors of no interest (three rigid-body translations and three rotations resulting from realignment). We used four contrasts: target-neutral, emotion-neutral, novel-neutral, and neutral-standard. In the “second-level” analysis, we averaged the four blocks of the task for each participant using FEAT with Fixed Effects. For estimation of group comparison (higher level, or “third level”) statistics, we used permutation testing utilising the FSL randomise tool, with threshold-free-cluster enhancement, which enhances cluster-like structures but remains fundamentally a voxel-wise statistical testing method ^39^. We used 5000 permutations and we report results at p=0.05 or less, family-wise error corrected for multiple comparisons, using the variance smoothing option (3mm) as recommended for experiments with small to modest sample sizes, as is common in fMRI research ^40^. For illustrative purposes only, we then extracted contrast values (contrast of parameter estimates, or COPEs in FSL) for each individual from voxels in which significant group differences were found (See bar chart in Figure 2B and 3B).

**Figure 2.**
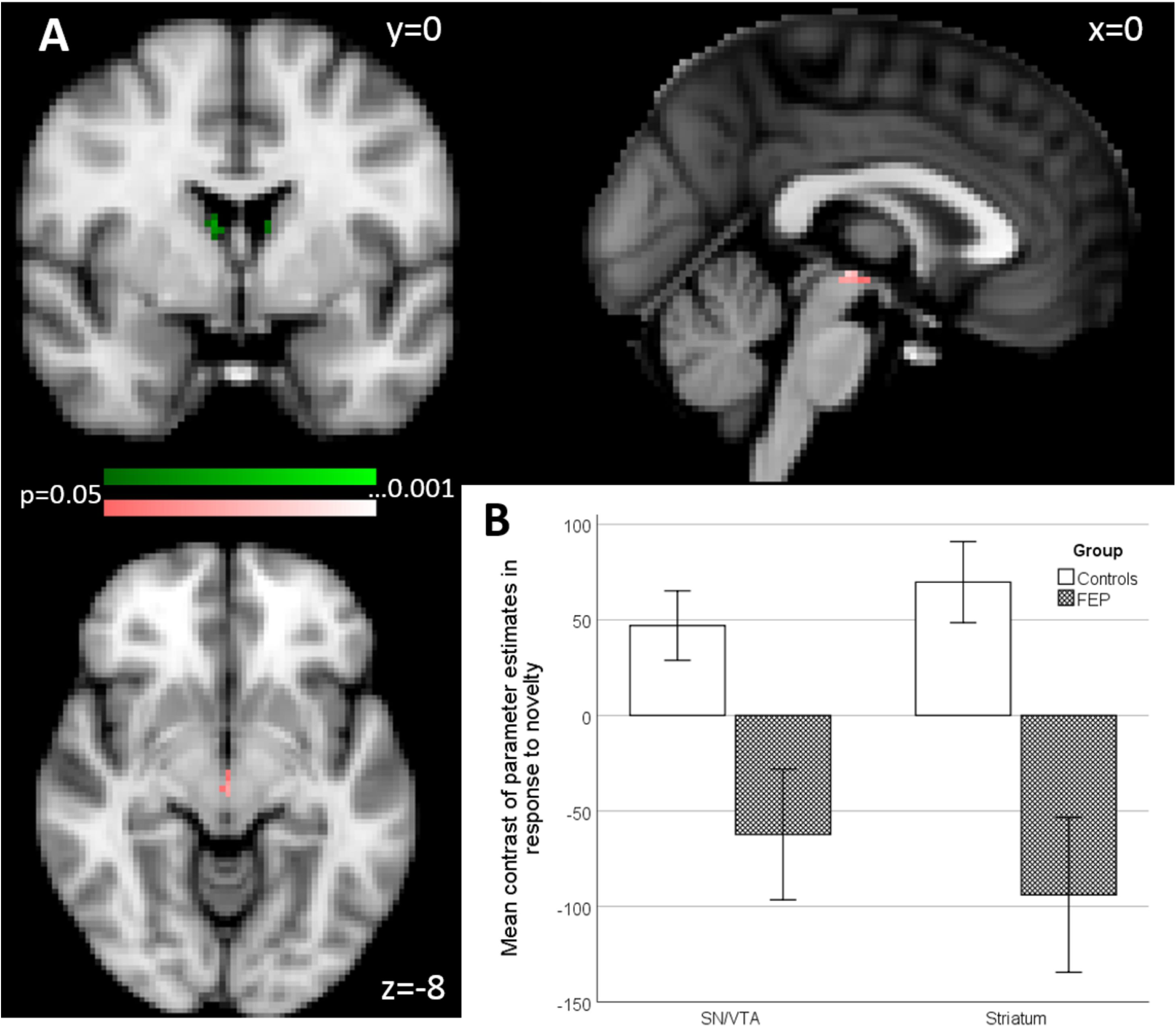
Group effects in primary and secondary region of interest (ROI) analysis of activation associated with novelty processing (novel oddballs versus neutral oddballs). A) Primary ROI (colour coding pink): SN/VTA, maximal difference at x=0, y=-20, z=-6. Secondary ROI (colour coding green), striatum, two clusters maximal difference at x=8, y=-2, z=14 and x=-8, y=-2, z=12 (p<0.05 FWE corrected). B) Bar chart shows the mean contrast (COPEs, FSL) values to group, extracted from significant clusters determined by FSL randomise ANOVA results of primary and secondary ROI analysis. Multiple significant clusters are combined. Error bars show ±1 SE.

Our main analysis was based on a region of interest (ROI) approach as follows. For all types of salience, our primary hypothesis involved the dopaminergic SN/VTA, which was used as our primary ROI. It was generated using the probabilistic atlas of Murty and colleagues ^41^, in which traditional anatomical segmentation was replicated using a seed-based functional connectivity approach and which provides a mask that consists of the SN and VTA, also used in our previous work ^22^. Furthermore, for each contrast we defined a secondary analysis, with either one region, or two non-adjacent regions combined in a single mask. For novelty, we used a secondary region of interest mask composed of the striatum (using a hand-drawn mask, encompassing both associative and limbic striatum, but not sensorimotor striatum, based on operational criteria ^40,41^) and the right lateral frontal cortex (utilising a sphere, 10mm, centred at x=50, y30=, z=28, based on our previous work ^21,22^). For negative emotional salience, our secondary region of interest mask was composed of the striatum and the amygdala (anatomically derived mask using the Harvard-Oxford subcortical structural atlas supplied with FSL). For targetness and rareness, we used the striatum as our secondary ROI.

Parameter estimates for the events that contribute to the contrasts of interest are presented in the supplementary materials (Supplementary Figures 5-8, and 11) for all conditions in primary and secondary ROIs. The parameter estimates indicate the potential drivers of the COPE (contrast of parameter estimates) effect.

For completeness, the same analysis as described above has been conducted on controls only and is presented in the supplements (Supplementary Figure 9, Supplementary Table 3 and 4).

**Table 3:**
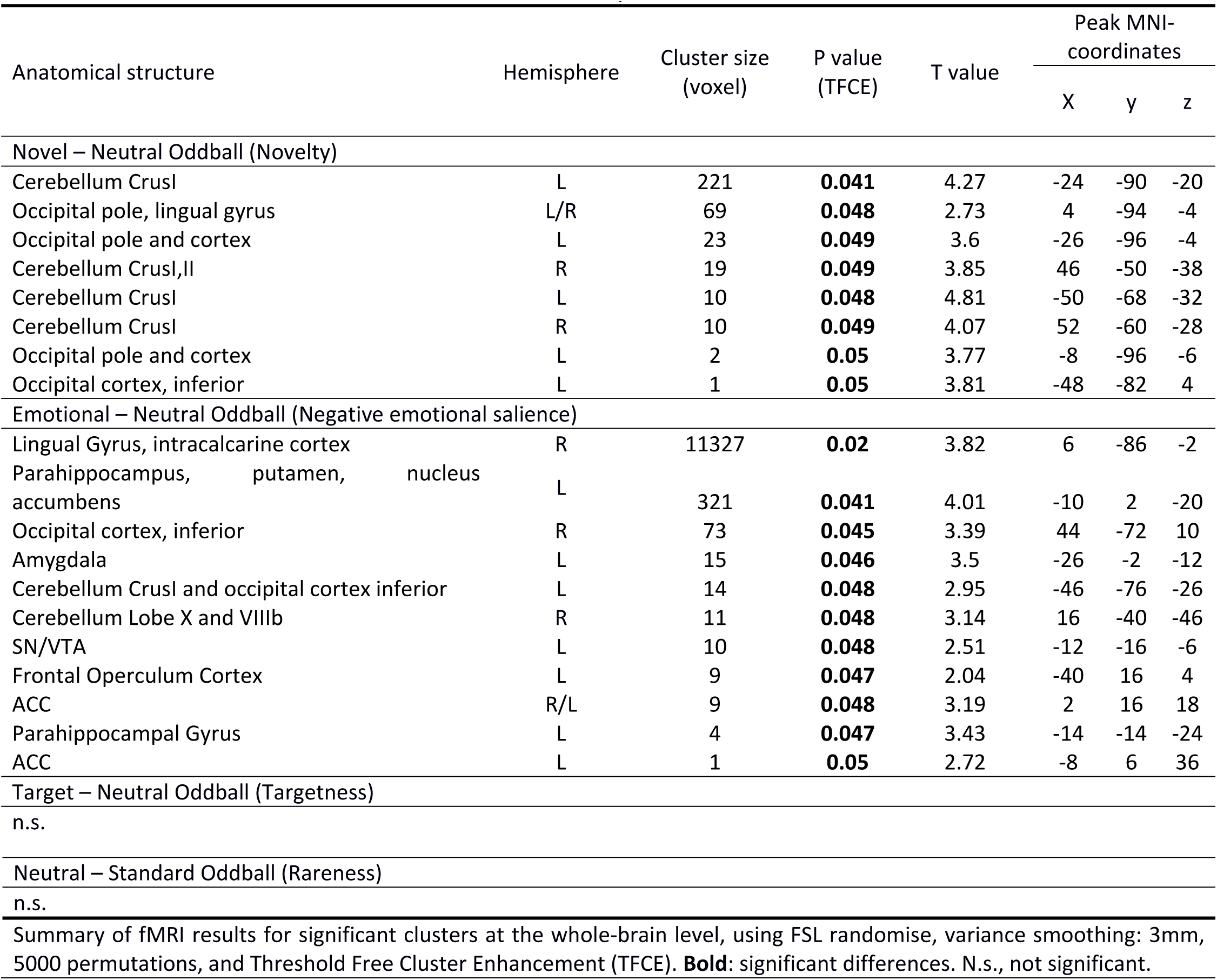
fMRI Activations from FSL randomise whole brain analysis.

### Movement differences during fMRI scan

As the task was relatively long (46min) and mostly passive (button presses were only required in 20 out of the total of 240 trials), we split the task into four blocks of 11.5min. Still many participants, independent of group, exhibited movements, possibly due to tiredness. We, therefore, excluded those blocks in which movement exceeded 3mm on average or 10mm maximum. In total, we identified 14 runs that fulfilled the movement exclusion criterion. Of those 14 runs, 10 were either from testing block 3 or 4, 2 were from testing block 2 and 2 were from testing block 1. Additionally, three runs had to be interrupted and were therefore not completed by the participants. If for a single participant only one or two runs remained for analysis, we excluded this participant entirely. Based on these criterions, we excluded three controls and one psychosis patient entirely, as well as one run in five psychosis patients and one run in three controls. We excluded those four individuals from all analyses within this study.

In the remaining sample, we compared the two groups in two separate repeated measure ANOVAs across the four testing blocks, one for movement means and one for maximum (Supplementary Figure 1 and 2). We did not find any significant group, run or interactions effect, neither for mean movement nor for maximum movement (all p>0.1).

## Results

### Demographic and questionnaire results

The demographic and rating results are summarised in Tables 1 and 2. There were significant differences in IQ between controls and patients (p=0.02). More importantly, however, the groups were matched in maternal education, which was similar across both groups (p=0.36). Alcohol consumption was significantly lower in psychosis patients compared to controls (p=0.002).

### Behavioural responses to pictures and reaction times

In order to maintain engagement with task, participants were required to press a button in response to the target picture. Due to technical problems, button presses were not recorded for eight controls, and one psychosis patient. Analysing the number of missed button presses and reaction times of the remaining participants across the four testing blocks (Supplementary Table 1), we did not find any significant effects for group, testing block or any interactions (all p>0.3). On average, participants missed to press the button on one target trial (mean: 1.0 SE±0.2) and generally required approximately 550ms (SE±0.02) to make a response, which is consistent with previous findings ^25^. Due to the high performance across all groups, we included the data of the nine participants without recorded button presses in all further analyses, in order to increase statistical power. We also repeated the analysis after excluding those participants. The results were very similar.

### fMRI results

*Novelty (novel-neutral oddballs)*

In our primary ROI, the SN/VTA, psychosis patients showed a significant reduction of activation compared to the controls (t=4.39, p=0.015 FWE corrected, 10 voxels; maximal difference at x=0, y=-20, z=-6). See Figure 2A and B.

In our secondary ROI, composed of the striatum and the DLPFC, we found two significant clusters, both within the striatum, that showed reduced activation for psychosis patients (cluster 1: t=4.51, p=0.03 FWE corrected, 18 voxels; maximal difference at x=8, y=-2, z=14; cluster 2: t=3.66, p=0.008 FWE corrected, 10 voxels; maximal difference at x=-8, y=-2, z=12). See Figure 2A and B.

On whole brain analysis using randomise, psychosis patients showed a significant reduction of activation in the occipital lobe, including the lingual gyrus and fusiform gyrus, and the cerebellum (Table 3 and Supplementary material).

### Negative emotional salience (emotion-neutral oddballs)

In our primary ROI, the SN/VTA, we found two clusters in which psychosis patients show significantly reduced activation(cluster 1: t=3.45, p=0.025 FWE corrected, 50 voxels; maximal difference at x=-12, y=-16, z=-6; cluster 2: t=4.37, p=0.02 FWE corrected, 22 voxels; maximal difference at x=0, y=-20, z=-4). See Figure 3A and B.

**Figure 3.**
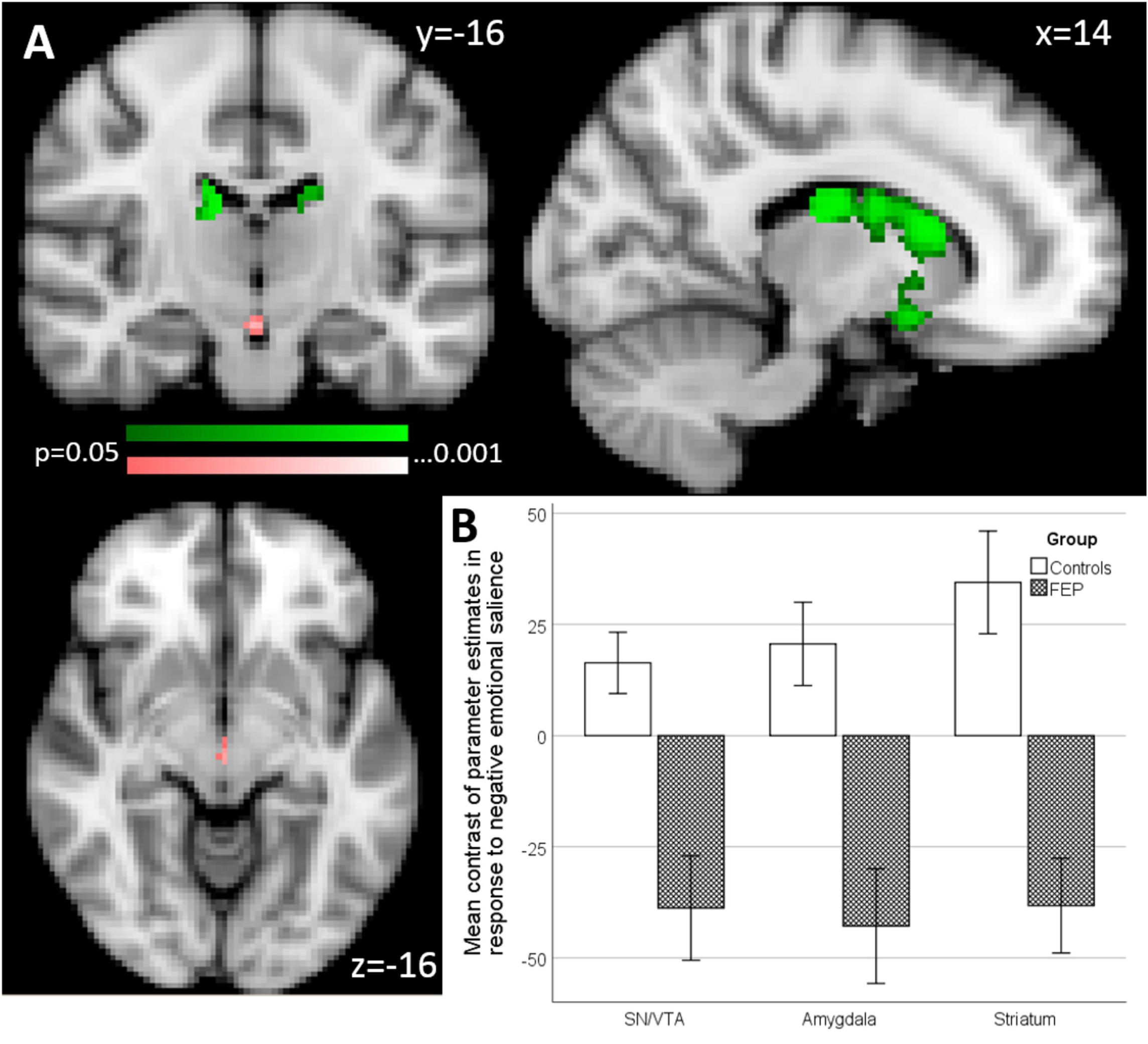
Group effects in primary and secondary region of interest (ROI) analysis of activation associated with negative emotional salience processing (emotional oddballs versus neutral oddballs). A) Primary ROI (colour coding pink): SN/VTA, four clusters maximal differences at x=-12, y=-16, z=-6; x=0, y=-20 z=-4; x=-10, y=-22, z=-20; and x=-12, y=-26, z=-20, and amygdala maximal differences at x=22, y=0, z=-14. Secondary ROI (colour coding green), striatum, four clusters, maximal differences at x=12, y=14, z=12; x=-10, y=-6, z=-16; x=-12, y=6, z=-14 and x=-14, y=22, z=-2 (p<0.05 FWE corrected). B) Bar chart shows the mean contrast (COPEs, FSL) values to group, extracted from significant clusters determined by FSL randomise ANOVA results of primary and secondary ROI analysis. Multiple significant clusters are combined. Error bars show ±1 SE.

In our secondary ROI, composed of the striatum and the amygdala, psychosis patients showed a significant reduction of activation compared to the controls in four clusters within the striatum (cluster 1: t=4.69, p=0.002 FWE corrected, 1141 voxels; maximal difference at x=12, y=14, z=12; cluster 2: t=4.11, p=0.008 FWE corrected, 320 voxels; maximal difference at x=-8, y=-6, z=-16; cluster 3: t=3.53, p=0.019 FWE corrected, 309 voxels; maximal difference at x=-12, y=6, z=-14; cluster 4: t=2.58, p=0.048 FWE corrected, 5 voxels; maximal difference at x=-14, y=22, z=-2). See Figure 3A and B.

On whole brain analysis, psychosis patients showed a significant reduction of activation in the amygdala, parahippocampal gyrus, lingual gyrus, striatum, cerebellum and the anterior cingulate gyrus (Table 3).

### Targetness (target-neutral oddballs)

In our primary ROI, the SN/VTA, psychosis patients showed a marginal reduction of activation compared to the controls (t=3.84, p=0.066 FWE corrected, 5 voxels; maximal difference at x=0, y=-22, z=-8).

On whole brain analysis, there were no significant group differences.

### Rareness/deviance (neutral oddballs-standard trials)

Our ROI analysis in the SN/VTA was not significant. Similarly, on the whole brain analysis, there were no group differences that passed our statistical threshold, corrected for multiple comparisons.

### Salience responses in controls only

The results for the same analysis as presented above have been conducted in controls only. Details for the whole brain analysis are presented in Supplementary Table 3, details for the ROI analysis in Supplementary Table 4. Furthermore, Supplementary Figure 9 shows the activation in the SN/VTA to emotional salience and novelty salience.

As novelty and negative emotional salience activate different, though adjacent, voxels also in the group comparison, we furthermore show the non-overlapping activation for the group comparison at various levels in the z direction (−6, −8, −10, −12, −14, −16,−18) in Supplementary Figure 10.

### Correlations of symptom score and brain responses in patients

We found positive correlations between SN/VTA signalling and the total score of negative symptoms (SANS; rho=0.66, p=0.047; Figure 4A), and delusions (P1; rho=0.77, p=0.002; Figure 4B) in response to novelty.

**Figure 4.**
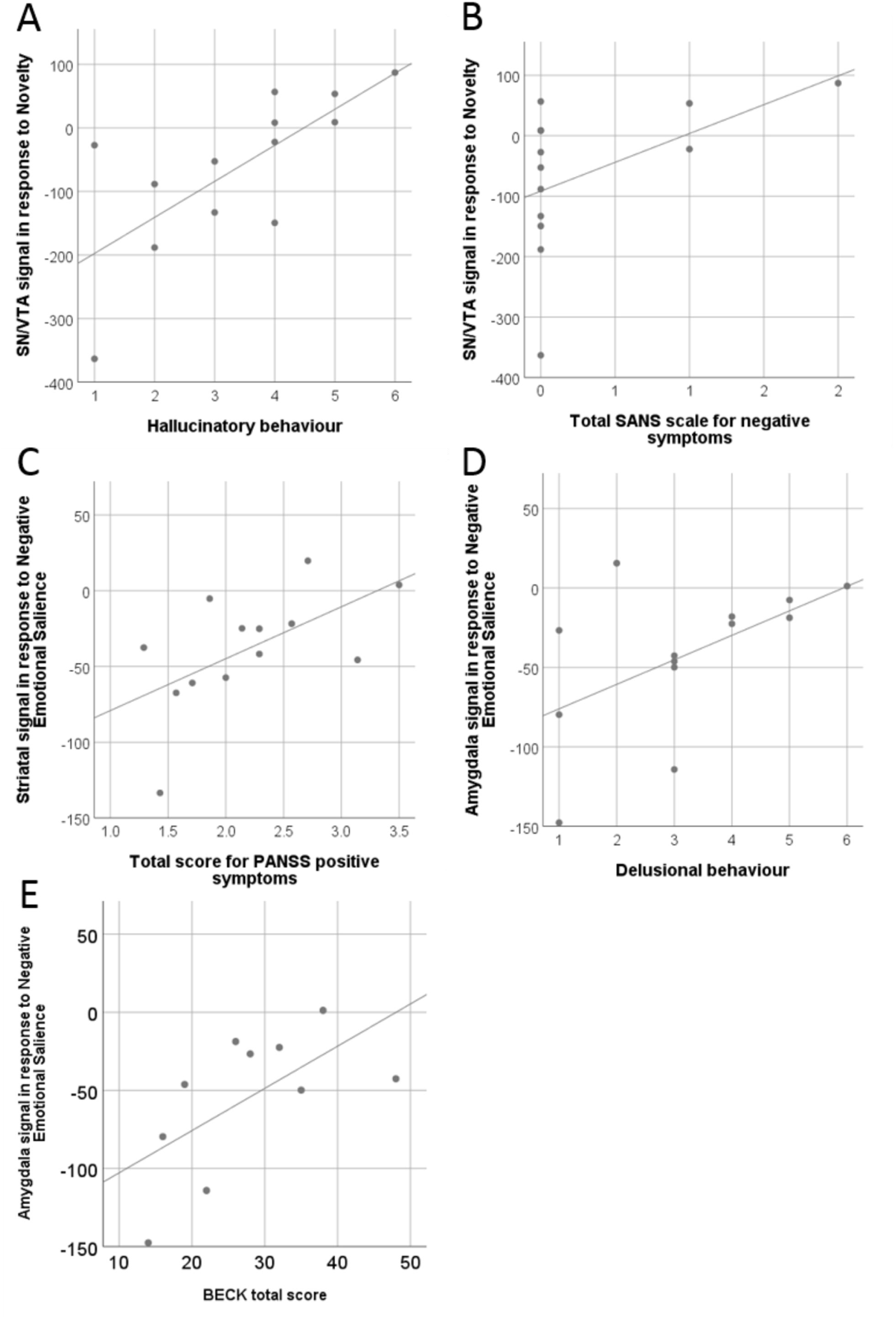
Correlation between signal strength and symptom score in patients. Panels A and B show significant symptom correlations with activation in the SN/VTA in response to novelty. Panel C shows significant correlation between total PANS score and striatal activation in response to negative emotional salience. Panel D and E show significant symptom correlations with activation in the amygdala in response to negative emotional salience. Lines indicate fitted regression lines.

Furthermore, in response to negative emotional salience, we found a positive correlations between striatal signalling and the total score of positive symptoms (PANS positive; rho=0.65, p=0.028; Figure 4C), a positive correlation between amygdala signalling and the Beck Depression Inventory (BDI; rho=0.64, p=0.048; Figure 4D), and a positive correlation between amygdala signalling and delusional behaviour (P3; rho=0.59, p=0.035; Figure 4E). Correlations were computed using a nonparametric Spearmen’s correlation.

We emphasize that we are presenting exploratory correlation analyses (given the small sample size); when controlling for multiple comparisons (correcting for all symptoms tested on each contrast) using a strict Bonferroni-approach only the correlation between SN/VTA signalling and delusional ideation in response to novelty would be retained as statistically significant. Additionally, we did not find any significant correlations between IQ and brain responses in any ROI (all p<0.05, no correction for multiple comparison applied).

## Discussion

We investigated brain responses to four different types of salience, including novelty, negative emotional salience, targetness and rareness/deviance in healthy volunteers and first episode psychosis patients. In psychosis patients, our results show reduced SN/VTA (primary ROI), striatal (secondary ROI) and cingulate (whole brain) signalling to novelty, reduced SN/VTA and amygdala (both primary ROIs), striatal (secondary ROI) and cingulate (whole brain) signalling to negative emotional salience, and reduced SN/VTA (primary ROI) and cingulate (whole brain) signalling to targetness. These results are unconfounded by antipsychotic medication as we tested an antipsychotic naïve patient sample. This study is first, to our knowledge, to present SN/VTA signalling alterations in psychosis in response to different forms of salience, all in absence of rewarding feedback. Our results, therefore, extend findings reporting midbrain and cortical abnormalities in response to reward prediction error signals in medicated and unmedicated psychosis patients at different stages of the disease ^19,21,42^, fronto-limbic connectivity alterations in response to novelty in acute, unmedicated psychosis patients ^43^ and striatal abnormalities in response to emotional salience in chronic, medicated and a small sample of unmedicated acute psychosis patients ^30^. Our results suggest that salience processing in the dopaminergic SN/VTA may be generally impaired in patients with psychosis.

The ‘aberrant salience’ hypothesis of psychosis postulates that dysregulated dopaminergic signalling in the mesolimbic system in people with psychosis results in the attribution of salience to irrelevant or non-significant stimuli ^10,44^. These unusually salient representations may lead to the formation of hallucinations or generally altered perceptions. As a result, patients may construct delusional explanations in order to explain these altered perceptions. Abnormal salience attribution is present from early and even prodromal stages of the disease ^19,23^. Usually, this theory is investigated in the context of motivational salience ^19,42^ using reward prediction paradims. Here, however, salience was investigated in the absence of the reward. For various forms of salience, the psychosis patient group showed significantly attenuated SN/VTA activation for the contrasts between salient and non-salient events. This impaired differentiation between salient and non-salient events could reflect dysregulated dopamine neuron activity, leading to the excessive attribution of salience to normally non-salient stimuli, and reducing salience to normally salient stimuli.

In healthy subjects, novelty identification is processed by a number of brain regions, including SN/VTA, striatum, parietal, and prefrontal cortices ^24,25,45^. Consistent with this, we observed group differences in the SN/VTA and the striatum in response to novelty. Our findings extend results of a study by Schott and colleagues ^43^; although, this study did not detect clear differences in the midbrain or striatum in psychosis, they found an increase in functional connectivity of the hippocampus and the orbitofrontal cortex with the rostral anterior cingulate gyrus and the ventral striatum. Our study also demonstrates significantly reduced activation in response to negative emotional salience compared to controls in right amygdala, the SN/VTA and the striatum in psychosis patients compared to controls. This result is consistent with the literature indicating reduced arousal to emotional stimuli ^46^. Our study also supports findings of a PET study indicating tonic over-activation of the amygdala and impaired striatal signalling during emotional salience processing ^30^. Jabbi and colleagues ^28^ reported increased dopaminergic releases in the amygdala and midbrain in response to emotional salience, which might be altered in psychosis. Our results, furthermore, reveal reduced activation in the thalamus of psychosis patients compared to healthy controls for negative emotional salience. The thalamus is a relay station of multiple neural connections and has dopaminergic synapses. Consistent with this and our findings, a study by Hadley and colleagues ^47^ reported reduced connectivity between the VTA/midbrain and the thalamus in schizophrenia patients.

In addition to reduced SN/VTA processing in response to novelty and negative emotional salience, we also found reduced signalling in response to targetness in patients. Therefore, our study is first to provide clear evidence for reduced SN/VTA processing in response to these different forms of non-motivational salience in psychosis. Together with the striatal findings of altered novelty and emotional salience signalling, the findings in the patients support the aberrant salience hypothesis for general (not just reward related) salience dysfunction. As both the midbrain and the striatum are dopaminergic key regions, it also provides supporting (though not definitive) evidence for a dysregulated dopaminergic system during salience processing in psychosis ^1^. In healthy controls, Bunzeck and Düzel ^25^ reported significantly enhanced SN/VTA activation in response to novelty, and also positive, but not statistically significant, activation in response to negative emotional salience, providing supportive evidence for a differential activation of the SN/VTA in response to novelty. Using a larger sample size than previous studies and a slightly different regional specification used for the SN/VTA, we, however, find significant SN/VTA activation to novelty and negative emotional salience in controls. Our results, therefore, support the view of general processing of salience in the SN/VTA ^7,8^, including novelty, negative emotional salience and targetness. An account reconciling these results with those of Bunzeck and Düzel ^25^, may be that SN/VTA is highly sensitive to novelty, but is also sensitive to (at least some) other forms of salience. On the other hand, we do note that the precise voxels within the SN/VTA that activate and/or show significant group differences to the different salient conditions do not overlap, though they are sometimes adjacent (see Supplementary Figure 9 and 10 for example). Furthermore, our focus of maximal group difference in novelty activation is slightly more rostral as compared to Bunzeck and Düzel’s findings of novelty associated activity in controls ^25^ but still lies within the SN/VTA ROI as defined by Murty et al.^41^. Given this variation in precise location of voxel clusters within the SN/VTA ROI, combined with the spatial resolution of the current study, we cannot definitively determine within this region whether the exact same neurons activate to diverse or specific stimuli ^48^. However, future studies could employ higher resolution at higher field strengths, to address these questions. Experiments using observational fMRI alone cannot prove that the abnormal SN/VTA patterns of various forms of salience signalling in psychosis have the same precise underlying pathophysiology, but future work combining the same or related fMRI paradigms with an intervention (pharmacological or brain stimulation) could further elucidate pathophysiological causal mechanisms.

Moreover, the whole brain analysis revealed reductions in anterior cingulate gyrus activity in psychosis patients compared to healthy controls in response to negative emotional salience. The cingulate cortex, as part of the salience network, has been found to show aberrant connectivity and structure in psychosis ^49,50^. We previously showed that the severity of psychotic symptoms in healthy volunteers induced by methamphetamine, significantly correlated with the degree of drug induced disruption of the incentive value signal disruption in the posterior cingulate cortex, suggesting a dopamine mediated mechanism in this region ^51^. A study by Gradin and colleagues ^52^ reported dysfunctional connectivity between the salience network and the midbrain during a reward learning task leading to abnormal reward processing in schizophrenia patients. Furthermore, structural alterations have consistently been documented in patients with psychosis^53–55^. Therefore, our results may provide an indication that possible dysfunctional interactions between the salience network and the SN/VTA may also lead to aberrant processing of different types of salience.

We expected to see increased activity in the visual cortices due to the use of a visual oddball paradigm across all stimuli ^45,56^. Here, we observed group differences in response to novelty and negative emotional salience, potentially in line with impaired visual perceptions often reported in schizophrenia (see review ^57^). In contrast with the previous literature, which reported hippocampal activity in response to salience ^25,43,58^, we did not find any activity in the hippocampus, neither in a group difference nor in a healthy volunteers separately. It is possible that signal in this region may not have been reliably captured during fMRI scanning.

In an exploratory analysis, we found positive correlations between SN/VTA activity to novelty and symptom scores for delusion and negative symptoms, between amygdala signalling to negative emotional salience and the Beck Depression Inventory and delusions, and between striatal signalling and total score for positive symptoms. However, when controlling for multiple comparisons, only the correlation between SN/VTA activation to novelty and delusions remains significant. Here, we would have rather predicted a negative correlation showing a decrease of SN/VTA activation with increased symptom scores, especially given the group difference that showed lower activation in the patient group as a whole. It is thought-provoking that in this small study, several forms of salience showed reduced activation across regions in the average patient, but greater activation associated with greater symptoms. One speculation is that reduced activation (between group results) could reflect a trait abnormality, and superimposed on this are state dysfunctions closely linked to symptom expression. However, symptom correlations with functional imaging have often yielded inconsistent results in schizophrenia research ^59^. One of the most important difficulties to reliably detect symptom correlations is gathering a large enough sample, and our small sample size of 14 patients is a clear limitation to assess symptom correlations. However, we report it to generate future hypotheses and to be available for future meta-analysis.

In conclusion, this study provides concise evidence for aberrant SN/VTA, striatal and cingulate signalling during non-motivational salience processing in a sample of antipsychotic naïve early psychosis patients. The results extend previous research by giving supportive evidence for the aberrant salience hypothesis of psychosis involving motivational and non-motivational forms of salience and the involvement of dopaminergic dysregulation in the development of psychotic disorder.

## Acknowledgements

The authors are grateful for the help of clinical staff in CAMEO for help with participant recruitment and Wolfson Brain Imaging Centre staff for MRI data collection support.

